# The predictive outfielder: a critical test across gravities

**DOI:** 10.1101/2024.01.08.574654

**Authors:** Borja Aguado, Joan López-Moliner

**Affiliations:** Vision and Control of Action (VISCA) Group, Department of Cognition, Development and Psychology of Education, Institut de Neurociències, Universitat de Barcelona, Passeig de la Vall d’Hebron 171, 08035 Barcelona, Catalonia, Spain; Sensorimotor Control and Learning group, Centre for Cognitive Science, Department of Human Sciences, Institute for Psychology / Centre for Cognitive Science, Technische Universitat Darmstadt, Germany

## Abstract

Intercepting moving targets, like fly balls, is a common challenge faced by several species. Historically, models attempting to explain this behavior in humans have relied on optical variables alone. Such models, while insightful, fall short in several respects, particularly in their lack of predictive capabilities. This absence of prediction limits the ability to plan movements or compensate for inherent sensorimotor delays. Moreover, these traditional models often imply that an outfielder must maintain a constant gaze on the target throughout to achieve successful interception. In this study, we present a new model that continuously updates its predictions, not just on the immediate trajectory of the ball, but also on its eventual landing position in the 3D scene and remaining flight time based on the outfielder’s real time movements. A distinct feature is the model’s adaptability to different gravitational scenarios, making its predictions inherently tailored to specific environmental conditions. By actively integrating gravity, our model produces trajectory predictions that can be validated against actual paths, providing a significant departure from previous models. To compare our model to the traditional ones, we conducted experiments within a virtual reality setting, strategically varying simulated gravity among other parameters. This gravity variation yielded qualitatively distinct predictions between error-nulling optical-based heuristics and our model. The trajectories, kinematic patterns and timing responses produced by participants were in good agreement with the predictions of our proposed model, suggesting a paradigm shift in our understanding of interceptive actions.

**Significance statement:** Catching a moving target, a challenge consistently faced across various species, exemplifies the complex interplay between perception, prediction, and motor action in dynamic environments. Prevailing models have been largely rooted in optical cues, often overlooking the predictive capacities essential for understanding real-world human behaviors and sidestepping crucial physical variables such as gravity. Our research introduces a novel model that emphasizes both the predictive component and the broader gravitational dynamics allowing for a more holistic understanding of interception tasks. This innovative approach not only holds implications for refining existing models of interception but also carries broader significance for training platforms, ensuring relevance across diverse settings, from Earth to altered gravity environments.

---

Consider how effortlessly an outfielder runs to catch a flyball. Explaining how this is achieved is the first step in addressing the general problem of interception, known as the outfielder problem (Chapman, 1968; Fajen and Warren, 2007; Michaels and Oudejans, 1992). The prevailing view is that the optical information obtained by our senses is sufficient to control an outfielder’s movements (Gibson, 1966), without delving into the prediction of future visual states. Consequently, attempted solutions have relied on heuristics defined by (a) maintaining a linear optical trajectory (McBeath et al., 1995) or (b) canceling the optical acceleration (Chapman, 1968; McLeod et al., 2006) while maintaining a constant velocity of the bearing angle (Fajen and Warren, 2007; Fink et al., 2009) to independently control the in-depth and lateral velocities, respectively (see Fig. 1 A).

**Fig. 1:**
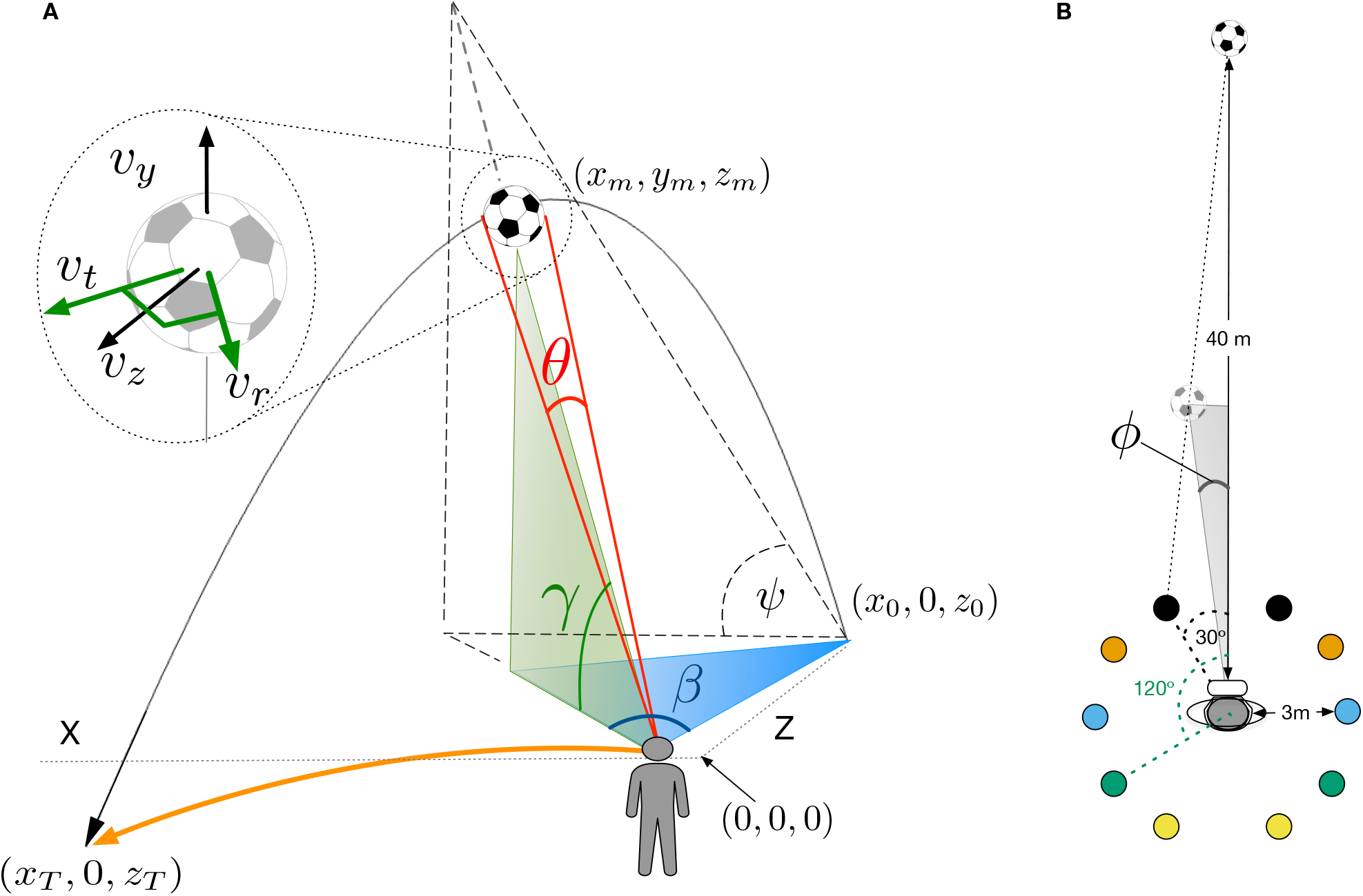
Main variables and trajectories: (A) Parabolic flight of a target and optical information available to an observer. The initial target position is (*x*_0_, 0, *z*_0_). Note that the origin is assumed to be at eye height. At time *m*, the ball is seen by the outfielder who has moved slightly from her initial position (0,0,0), that is, the origin. At this time, the ball subtends a retinal size (*θ*) and elevation angle (*γ*). *β* (azimuth) is the angle between the projection of the ball on the floor, observer, and initial (home) position. *Ψ* denotes the optical trajectory projection, where tan *Ψ* = tan *γ*/ tan *β*. The inset shows different velocity vectors for the movement of the ball. The radial component *υ* _*r*_ is the component in the direction of the observer, which is orthogonal to the tangential component *υ*_*t*_ (green color). The vertical component *υ*_*y*_ and the depth component *υ*_*z*_ are shown in black. (B) Different trajectories used in our experiment (top view). The initial position of the observer represented the origin. Colored circles denote the final landing position at 3 m from the observer, at 30, 60, 90, 120 and 150 degs relative to the observer’s initial position (only two angle values are shown). Ball’s initial position was set 40m away from the actor. *ϕ* denotes the bearing angle (the angle between the observer’s heading and current ball position). The GOAC model proposes that the acceleration of tan *γ* is nullified and the velocity of tan *ϕ* would be roughly constant to control in-depth and lateral movement, respectively McLeod et al. (2006). Keeping the change of *Ψ* linear represents the heuristic proposed by the LOT model McBeath et al. (1995).

The role of prediction in ball catching was highlighted in early studies (Sharp and Whiting, 1974) to address uncertainty due to occlusions and sensorimotor delays (Nijhawan, 1994). However, such a perspective faces skepticism (Zhao and Warren, 2014) due to the perceived complexity of predictive models. Models anticipating ball trajectories based on initial conditions (Saxberg, 1987) have been deemed unlikely due to the intricate internal models needed to anticipate projectile motion (Fink et al., 2009; McBeath et al., 1995; Shaffer and McBeath, 2005). Yet, compelling evidence emerges from cricket, where predictive saccades reveal anticipatory eye movements that precede the actual ball trajectory (Land and McLeod, 2000), and adapt to the elasticity of the ball at play (Diaz et al., 2013). In this line, (de la Malla and López-Moliner, 2015) shows how initial information seamlessly integrates with late-stage data, suggesting a continuous updating of prediction. These findings bridge sensory online cues with anticipatory strategies, offering a more intricate view of dynamic interception tasks.

Despite their fundamental differences, all of these models predict successful catches, that is, the same ending positions with little difference between the traveled paths of an outfielder. This introduces further challenges when testing different models against the experimental data. Yet, one constant that remains absent but significantly impacts the optical variables’ trajectory is Earth’s gravitational acceleration despite Gibson (Gibson, 1979) acknowledging gravity’s key role as an environmental reference. Several studies suggest that gravity is not only employed when catching objects in free-fall (McIntyre et al., 2001; Zago et al., 2009) but that humans inherently leverage Earth’s gravity as a useful prior for predicting motion (Jörges and López-Moliner, 2017).

Here, we demonstrate that when monocular optical variables are combined into a model with known constants, such as gravity and physical size, it is possible to accurately predict when and where a ball will land relative to an actor in the general case. Furthermore, the same model can be used to update these predictions to guide the outfielder to the landing position by integrating the ongoing predictions into a simple controller. An experiment using immersive virtual reality (see Fig. 1 and Methods) enabled us to manipulate the ball size and gravitation, introducing values that deviated from the expected ones. Assuming that people use Earth’s gravity as a prior (McIntyre et al., 2001), the model predicts specific paths for the different simulated gravity values, which are qualitatively distinguishable from paths predicted by previous models using optical variables alone in the same conditions. The proposed model can predict the empirical trajectories and provide a very good account of the observed actor’s kinematics, an aspect that some previous models (see Jacobs et al. (1996)) have failed to predict.

## Results

The data and R code to run the analysis, simulations, obtain predictions for the different models, and reproduce the main figures of the article is available in the following OSF project (link). An interactive R-based shiny application (4_Predictive_Actor.R) allows for the exploration of different model parameters and the resulting trajectories of the simulated actor.

### Gravity affects spatial trajectories

In our experiment, participants had to run towards where they thought the ball would land (see Fig. 1 B for different trajectory angles). They pressed a button when they considered the ball was again at eye height (initial vertical position) after launch without seeing the final 10% of the trajectory due to occlusion. To study the locomotion towards the interception location, we computed the average heading, defined as the angle between locomotion direction and the ball’s initial position. We did so under different conditions defined by trajectory angle, gravitation, and ball size (see Methods for further details). Fig. 2 A shows this average direction of the spatial trajectory across participants for different simulated gravities. Individual trajectories are shown in Fig. S1 in the Supplementary Information (SI). In agreement with previous studies (Fink et al., 2009; McBeath et al., 1995), participants did not follow a linear path. Instead, when the ball landed in front of them, they followed a slightly convex trajectory, and when the ball landed behind them, their path was concave. Importantly, when analyzing the segment of the trajectory before occlusion and during the actor’s movement, gravitation had a significant effect on heading angle (Fig. 2 A, F(2,20) = 64.42, p < .001, *η*^2^ = 0.19). When gravity exceeded terrestrial levels, participants’ paths deviated more from the ball’s trajectory: becoming more convex when running forward and less concave when moving backward.

**Fig. 2:**
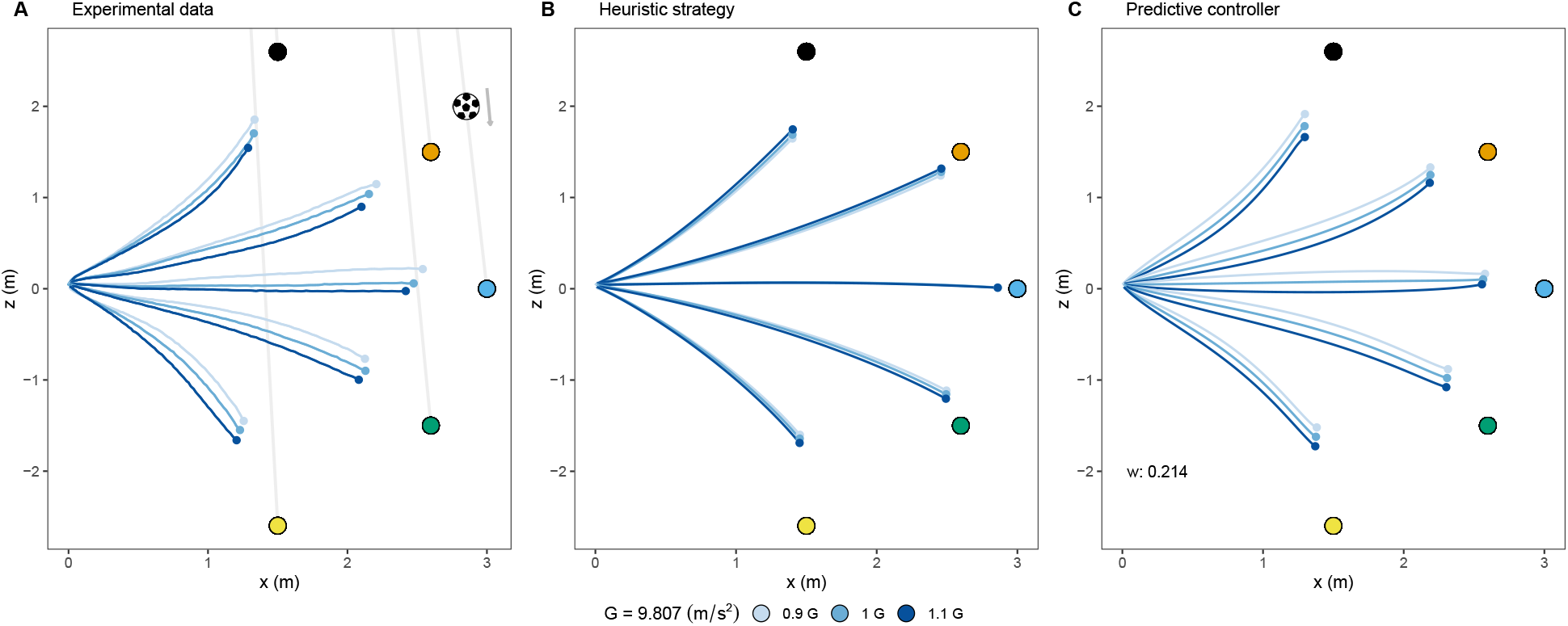
Empirical and simulated paths: (A) Average empirical paths followed by our participants in the experiment. Gray lines indicate the trajectories followed by the ball. (B) Paths simulated using a controller guided by GOAC heuristic strategy. (C) Paths simulated using a controller guided by a predictive strategy. The weight of the terrestrial gravitation prior that provided the best fit was 0.192.

Since gravity, flight duration, and initial vertical velocity 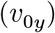 are intrinsically related, we needed to ensure that gravity is the primary variable influencing trajectory differences, not 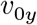or flight duration. Our experiment introduced variability in flight duration by varying 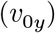, enabling us to examine its impact. In the Supplementary Information (Fig. S2A), we demonstrate that variations in 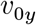 do not account for the observed differences in trajectories landing in front of the observer, as confirmed by the inset showing no correlation between 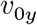 and in-depth landing position. This analysis points to gravity as the key explanatory variable.

We compared empirical data against predictions from various strategies, including an actor implementing the GOAC heuristic, which controls radial and tangential velocity by nulling the acceleration of tan(*γ*) and the velocity of tan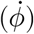, respectively (details in Methods). Fig. 2 B shows predicted paths using the GOAC strategy, indicating that despite its ability to replicate curved paths, it does not match the empirical differences across gravities observed in front of the actor: larger gravities yield less convex paths.

Moreover, an actor based on our model’s predictions aligns more closely with the empirical data, as seen in Fig. 2 C. This model replicates the curved paths and heading directions across different gravity values, with larger gravity leading to more convex forward paths and less concavity when moving backward. This alignment is further supported by cross-correlation analyses (Fig. S2A), showing a stronger correlation between heading and our model’s control variables compared to those of heuristic strategies. Our model combines Earth’s gravitational acceleration (*g*_*e*_) and simulated gravity (*g*_*s*_) with weights of 0.192 and 0.808, respectively, for optimal empirical fit (Fig. S3, SI). The effective gravity (*g*) in the model equations is *g* = *wg*_*e*_ + (1 − *w*)*g*_*s*_. Crucially, participants can discern *g*_*s*_ through changes in specific optical variables, integrated into our model. The inset of Fig. S2B and Fig. S3 illustrate how these variables provide sensory cues for gravity perception and the impact of varying weights on spatial trajectory separation under different gravity conditions.

While the initial impression might suggest that our fitting procedure favors our predictive model primarily due to the additional parameter (w), it is essential to highlight a pivotal observation from Fig. 2: the symmetry observed in the heuristic controller’s predictions for different gravities, both for trajectories landing in front and behind. This symmetry emerges from the principles of the GOAC model and underscores a qualitative difference between the heuristic controller’s predictions and the empirical data. To investigate whether this qualitative discrepancy can be rectified by introducing more flexibility into the heuristic controller, we conducted additional fittings. In these fittings, we granted the heuristic controller more flexibility by freeing one controller term which consist of two parameters (see Fig. S4 in the SI) and subsequently two controller terms (see Fig. S5 and Table S1). Despite this heightened flexibility, the heuristic controller remains incapable of aligning its predictions with the nuanced trajectory trends observed in the data. These tests, then, demonstrate that the qualitative disparity observed between the heuristic controller’s predictions and the empirical data, cannot be overcome to reverse the predicted paths to match the observed ones. This reaffirms our confidence in the predictive model’s superior fit and its ability to faithfully represent the empirical data.

While the simulated gravitational acceleration had a pronounced impact, the ball size influenced trajectories only marginally (F(2,20) = 2.99, p = .083, *η*^2^ = 0.005). Our model predicts an effect of size, specifically if participants use the mean size as a prior due to familiarity. Though a one-tailed test could be considered based on this prediction, the modest effect size warrants a cautious interpretation, suggesting reliance on the actual size without drawing on prior knowledge. This perspective is expanded upon in the Discussion. There was no significant interaction between the simulated gravitational acceleration and ball size (F(4,40) = 1.02, p = .402, *η*^2^= 0.003).

### Predictive model

Unlike previous models in which the outfielder movement is coupled with the ball through an error-nulling tactic, our actor uses optical variables combined with knowledge about physical size and gravity to obtain first an estimate of the remaining flight time *T*_*c*_ or time-to-contact, TTC (see the model section for a complete derivation):

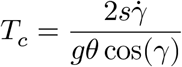

In this equation, *s* represents the ball’s physical size, and *g* is gravitational acceleration, as previously discussed, influenced by weights (*w* and 1 *− w*) assigned to terrestrial (1G) and simulated gravitational acceleration. Both *γ* (elevation angle) and *θ* (ball’s angular size) are time-varying factors, as illustrated in Fig. 1 A. We have omitted the time indices from our descriptions for clarity. At the trajectory’s onset (i.e., *t* = 0), given that *γ* = 0 and *cos*(0) = 1, the equation simplifies to 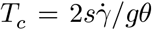. It is crucial to emphasize the role of the observer’s movement in this context. The equation offers an estimate of *T*_*c*_ for a specific system state. This state is dynamic and is influenced by how the observer moves. Specifically, angular variables *γ* and *θ* are directly affected by the observer’s movement. As such, the way in which participants move and adjust their position plays a crucial role in the real-time calculations of *T*_*c*_. Importantly, this initial temporal estimate quickly allows the actor to obtain an accurate estimate of the final lateral *x*_*T*_ position of the ball relative to her, thus making spatial and temporal information inseparable (Tresilian, 1999):

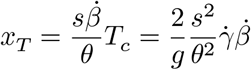

where 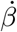 is the rate of change of the azimuth angle (see Fig. 1 A) and the final position *z*_*T*_ in depth relative to the observer:

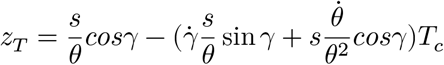

It is important to note that the reference frame in which *x*_*T*_ and *z*_*T*_ are updated is not fixed; it rotates as the observer turns. Fig. 3 shows the time course of these three estimates (*T*_*c*_, *x*_*T*_ and *z*_*T*_) for three of the trajectory angles, considering the changing position of the actor under terrestrial acceleration. In other words, Fig. 3 tells us how the temporal and spatial predictions are updated. The data presented in this figure are representative of trials in which the participants were able to catch the ball. We classified a trial as a catch only if the observer was within 0.5 meters of the ending point at the time of contact. We plot both estimates based on our actors’ actual movements (dots) and the resulting best fit (line) from the predictive controller (see next section). Notably, while the actual paths in space were curved, participants took a route where timely changing positions resulted in a linear decrease in the remaining flight time estimate, *T*_*c*_, as shown in Fig. 3 A. When an actor’s movements linearize *T*_*c*_, it consistently reduces the estimated final lateral position according to the previous equations (Fig. 3 B). Finally, Fig. 3 C shows the corresponding estimate for the final depth position of the ball relative to that of the observer. The deviations of data points from the line in Fig. 3 stem from variations in actor positions at the same time frames for these trajectory angles, influencing the updated estimates. The increased fluctuation observed in Fig. 3 C is a result of noisier estimates when updating *z*_*T*_ due to the consideration of retinal expansion of the ball (see previous equation). The accuracy of these estimates depends on both the actor’s position and the timing of these positions. At the flight’s onset, these estimates accurately reflect the relevant physical aspects in the 3D environment (e.g. landing positions and flight time), regardless of the observer’s initial position relative to the ball; but accuracy of later estimates will depend on the actual movement of the actor. The model can provide sufficient information to guide the movement to obtain a linear decrease in the remaining flight time. Fig. S6 in the SI shows an error map of the temporal estimates for all possible positions at different times. An animated version of this figure (“temporal_error_map.gif”) is also available in the SI. This plot shows that, irrespective of the position of the observer at the target launch, the error of temporal estimates is minimal at the beginning.

**Fig. 3:**
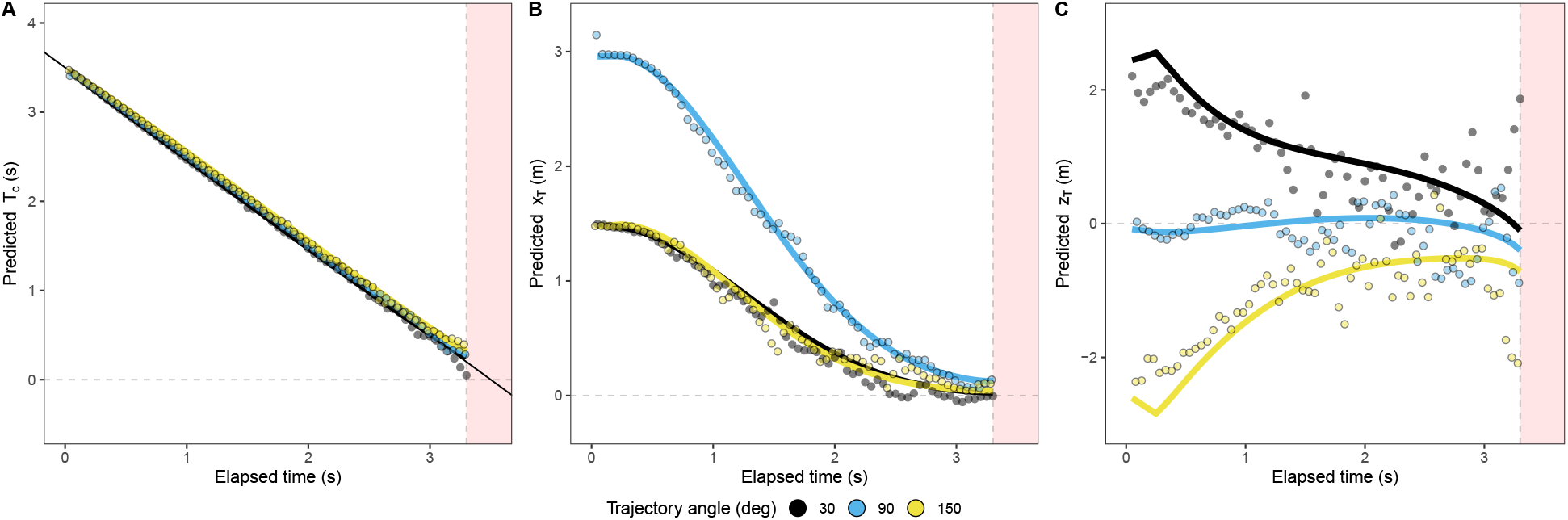
Temporal course model estimates: Panels (A), (B), and (C) show estimates based on our predictive model for successful catches: remaining time-to-contact (*T*_*c*_) and distance to the ending position in the x (*x*_*T*_) and z-axis (*z*_*T*_) respectively. The estimates based on the actual movements are represented by points, and the best fit of the model controller across all the trials is represented by a line. The color code represents three of the trajectory angles tested in the experiment. The red area indicates the occlusion interval.

While Fig. 3 illustrates the dynamic spatio-temporal estimates generated by our model, Fig. S7 in the SI presents the optical variables associated with heuristic strategies, which are derived from the same movements as in Fig. 3. These optical variables align with the expected patterns predicted by their respective strategies, allowing the interpretation that these heuristics are a result of employing optimal strategies, as previously suggested by (Belousov et al., 2016).

### The controller

To generate the trajectories based on the spatio-temporal estimates shown in Fig. 3 and make the predictions plotted in Fig. 3 C, we implemented a simple dynamical model in which an actor controls the radial velocity *(υ* _*r*_, movement towards the ball) and tangential speed (*υ*_*t*_, orthogonal component to *υ*_*r*_) of her movement independently (see controller dynamics in the Methods). The actor was required to wait for 350 ms before initiating movement, which included a simulated processing sensorimotor delay of 200 ms (Brenner and Smeets, 2015).

At each time, the estimates of the two velocity components were computed as *υ* _*r*_ = *z*_*T*_ /*T*_*c*_ and *υ* _*t*_ = *x*_*T*_ /*T*_*c*_, and an acceleration component was computed in the dynamical model and integrated to keep the velocity of the actor close to the estimates. Our model estimates provide the actor knowledge of the average velocity at which she must move laterally and in depth to reach the final position of the ball. This strategy appears intuitive and is consistent with the subjective feeling of quickly knowing whether one would be able to catch the ball (Postma et al., 2018) and decide to start running. Notably, we can provide, as discussed later, a testable approach to assess catchability based on the model’s estimates. However, this does not necessarily imply that people can explicitly tell where the ball will land (Shaffer and McBeath, 2005).

### Movement kinematics

The predictive model, based on the controller we have just presented, effectively accounts for the observed kinematics. In the first column of Fig. 4, we display the average lateral velocity component (top panels) and the depth velocity component (bottom panels) across participants, encompassing various trajectory angles and gravitational conditions. The second and third columns showcase the predictions for the GOAC strategy and our predictive strategy, respectively. Our predictive model exhibits a notably better fit to the kinematic data (***R***^2^=0.98, lateral; ***R***^2^=0.983, in-depth) compared to the heuristic counterpart (***R***^2^=0.94, lateral; ***R***^2^=0.96, in-depth). The GOAC model fails to capture the observed lateral deceleration during the latter portion of the trajectory, typically occurring after 1.5 seconds. This limitation in replicating the deceleration aligns with a known shortcoming of the LOT model too, as previously discussed in (Jacobs et al., 1996). Importantly, our model is the only one that demonstrates some sensitivity to gravitational variations in the depth component of movement, where gravity exerts a substantial influence (F(2,20) = 35.22, p < .001, *η*^2^ = 0.058) considering the whole trajectory. In this dimension, we observe a significant impact of gravity on the trajectories. However, our model falls short in capturing the variations in lateral movement, where gravity still has a significant effect (F(2,20) = 8.35, p = .02, *η*^2^ = 0.018).

**Fig. 4:**
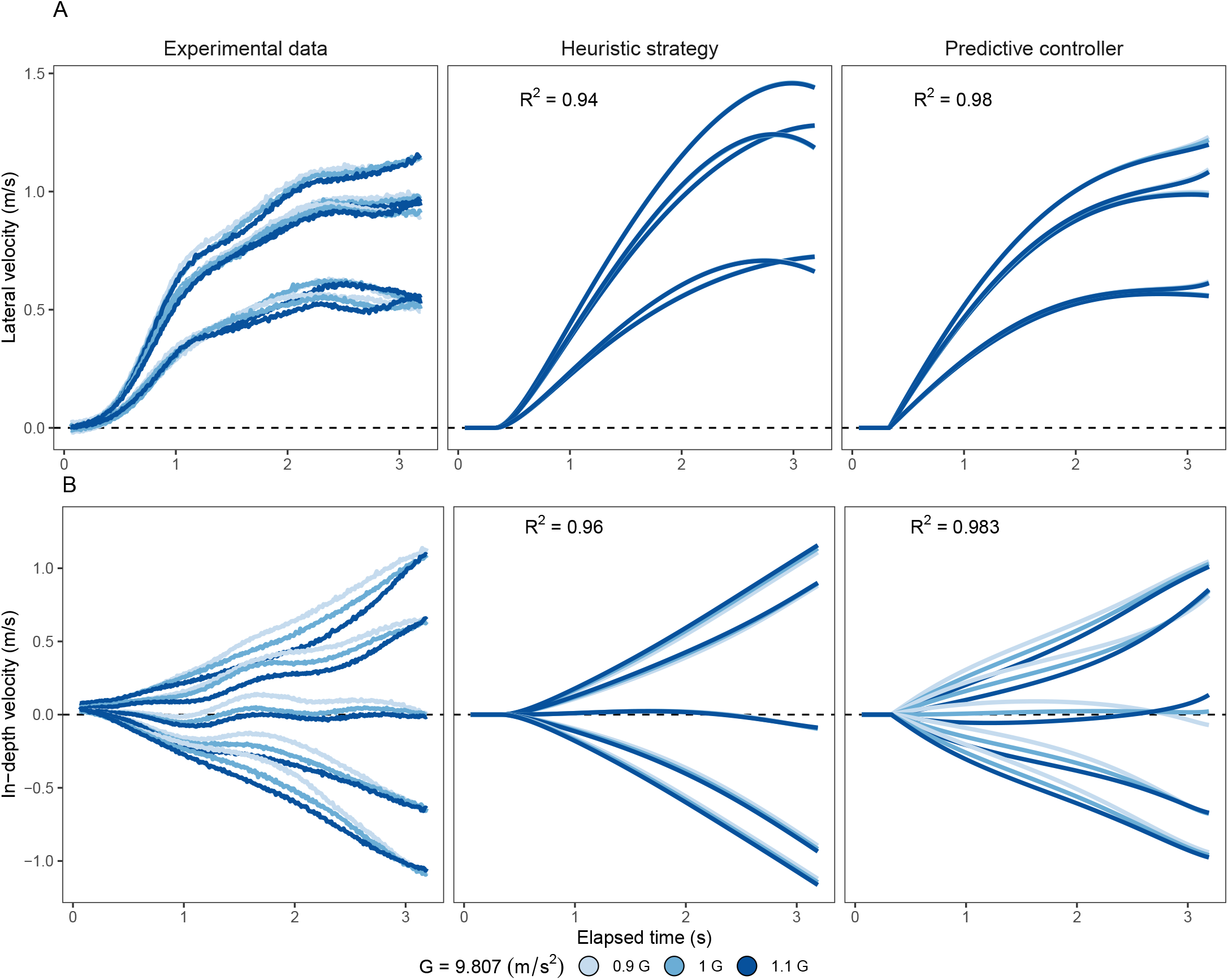
Movement kinematics: Average velocity for the lateral (top panels) and in-depth (bottom panels) motion across trajectories and gravitations (color code). The columns present experimental data, heuristic predictions, and predictive model predictions. Experimental data was smoothed by using a moving average (window of 0.1 s) that was run in the two directions to prevent a phase shift.

### Final stage: temporal judgments

According to our model predictions, the assumption of terrestrial gravitation to some extent will lead to an overestimation of time-to-contact (TTC) when an actor is exposed to gravitations larger than terrestrial (1G). Indeed, this is the pattern we observed (Fig. 5 A; Gravity-main effect: *F*(2, 20) = 33.802, *p* < .001, *η*^2^ =0.004). The slopes in Fig. 5 A predict the trend of the expected response pattern if Earth gravity was given all the weight (*w* = 1) and weaker weights (*w* = .192 and *w* = .5). The mean weight for Earth gravity that accounts for the separation of the spatial trajectories across gravitational accelerations (w=0.192) is smaller than the weight explaining the effects of gravity shown in Fig. 5 A. The 95%-CI of the fitted slope across temporal errors does not include the weight of 0.192 across Gravitations (see shaded area in Fig. 5 A). The assumption of a soccer ball size as a prior would lead to underestimation of TTC if the ball size is larger than expected and vice versa. This is the trend shown by the data points in Fig. 5 B. Participants underestimated the TTC when balls of a larger size were present and vice versa (Size-main effect: *F*(2, 20) = 25.268, *p* < .001, *η*^2^=0.005). Note that none of the heuristic strategies make different predictions for different sizes with respect to the final response time. Gravitation and ball size were not the only potential factors influencing the estimates of the remaining TTC. Since the accuracy of the estimates relies on the timing of the observer positions, and they did not proceed to the final landing position (see Fig. 2 A), the final estimates of TTC may not be perfectly accurate. This inaccuracy depends on whether the ball falls in front of or behind the observer’s line of sight (Aguado and López-Moliner (2021) and inset of Fig. 5 C). We can exploit this fact to see if the observed TTC estimates for the different trajectories are consistent with the predicted biases shown in the inset. Fig. 5 C shows that this is certainly the case with the pattern of errors found in our results (initial angle main effect: *F*(4, 40) = 7.239, *p* < .001, *η*^2^=0.012). Note that the direction of the biases in the TTC estimates is predicted by the final available information from the model depending on the actual position of the actor. This provides strong evidence for the use of predictive information from the proposed model. This additional evidence, combined with previously reported findings such as heading and kinematics, further solidifies our model as the most comprehensive explanation for all observed data patterns.

**Fig. 5:**
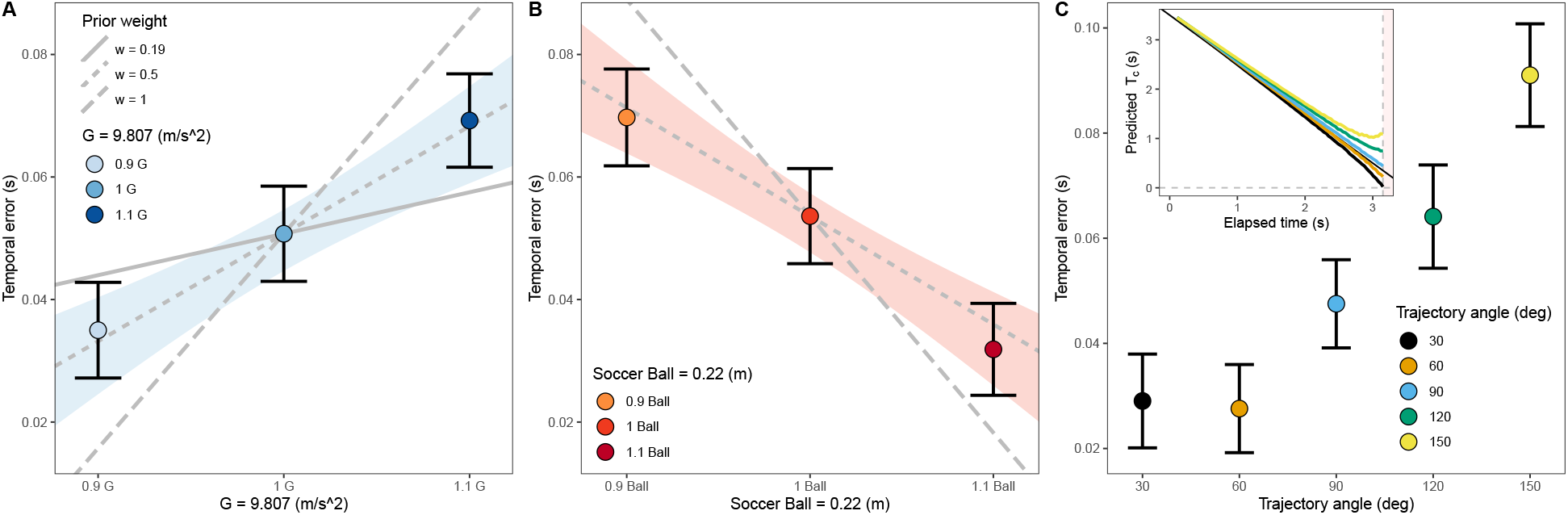
Temporal error per gravity, size and trajectory angle: Averaged temporal errors committed by our participants across gravitations (A) ball sizes (B) and trajectory angles (C) present in the experiment. Error bars indicate 95% confidence interval. Lines in panels (A) and (B) indicate predicted temporal errors, assuming a combination of prior knowledge (w) and veridical physical variables (1-w) for Gravity in panel A and size in panel B. The shaded area indicates a 95% confidence interval for the fitted linear models that only includes the slope resulting from a prior weight of a 0.5 for Gravity and Size. In panel (C), the inset represents the average predicted time-to-contact per trajectory derived from our participants action.

## Discussion

Dominant theories explaining how humans catch parabolic balls rely on error-nulling tactics (McBeath et al., 1995; McLeod et al., 2006). These theories suggest that outfielder movements are linked to optical variables related to the ball’s motion. While this coupling always predicts successful catches, it can be challenging to identify the specific controlling variable unless perturbed, non-parabolic trajectories, are shown (Fink et al., 2009). We propose a predictive model that combines optical variables and two physical constants that are assumed to be known to the participant, gravity and object size, which play no role in previous heuristic strategies. Our model predicts systematic errors when gravitation or ball size deviates from the value assumed by the observer. Therefore, we experimentally manipulated gravitation and ball size to investigate whether our participants’ behavior was consistent with the deviations predicted by our model or with previous heuristics in two different stages: locomotion and final temporal judgement. Converging evidence from locomotion, kinematics, and timing responses biases establishes our model as the most comprehensive explanation of the data. Our participants took different paths towards the interception location depending on the exposed gravitational acceleration during the locomotion phase. These paths would be consistent with an incorrect extrapolation of the ball’s motion due to a misestimation of the remaining TTC caused by assuming an Earth’s gravity prior to some extent. When the model predicts an underestimation of the TTC, the observer moves in the direction of the ball as if it falls short and vice versa. This is exactly the pattern found in our experiment (see Fig. 2 A). Importantly, note the opposite predictions for some of the trajectories between our model and heuristic strategies. For example, when gravitations larger than terrestrial are simulated, coupling the actor movements with an optical variable such as the elevation angle will predict movements slightly forward towards the ball (see Fig. 2 B), because a larger gravity will pull the ball down to a greater extent, reducing the elevation angle (*γ*). We observe the opposite pattern, which is consistent with weighting Earth gravity in our model, resulting in a smaller perceived value of gravitational acceleration than that actually shown. This results in an overestimation of TTC, causing the actor to move away from the ball. Qualitatively distinct predictions from the heuristic strategy remain unchanged even when introducing more free parameters into the heuristic controller (as evidenced in SI Figs. S4 and S5). Consequently, it becomes challenging for various heuristic strategies to account for these seemingly arbitrary trajectories. The previous study by Fink and colleagues (Fink et al., 2009) stands out as one of the few studies that contrasts the GOAC and LOT models in ball-catching scenarios, finding evidence for the GOAC model. Although this might appear contradictory to our main findings, it is important to note that under conditions without gravity manipulation, such as in Fink’s study, our model’s predictions closely align with GOAC, to the point of being nearly indistinguishable (as evident in our SI figures). The distinction arises only when manipulating gravity.

Unlike gravity, size manipulation only had a marginal effect on the trajectories. This result is consistent with participants giving very little weight to a prior size or accessing the correct size of the ball while it was visible. This can be explained by the use of available binocular information (Berkeley, 1709; Regan et al., 1979), by (correctly) assuming a constant initial distance (Hecht et al., 1996; McConnell et al., 1998; Watson et al., 1992), or using a combination of both sources of information. However, like gravity, prior size due to familiarity (Hosking and Crassini, 2011; López-Moliner et al., 2007) did have an effect on the temporal response (second stage), which was performed after occlusion, which is consistent with prior information becoming more relevant when sensory evidence is absent (Körding and Wolpert, 2004). Prior of known size would become more relevant in the second stage (e.g., judging the remaining TTC or adjusting the final catch) than for guiding locomotion.

Emphasizing consistency with prediction, our results reveal that participants in-depth speeds of various trajectories separate quite early on, within less than 0.5 seconds (see Fig S8 in SI). This happens even when changes in the elevation angle remain below known acceleration detection thresholds (Gottsdanker et al., 1961; Werkhoven et al., 1992). The observed separation aligns with our estimates used as control variables (also in Fig S8) where differences between estimates are well above threshold values. Previous predictive solutions (Tresilian, 1995) assumed that the ideal path towards the interception location would be straight. However, our experimental results show that observers consistently follow a slightly curved path towards the interception location, which is in agreement with previous studies (Fink et al., 2009; McBeath et al., 1995; Mcleod et al., 2001). Our kinematic analysis (Fig. 4 A, B) reveals that actors initially prefer lateral movement before moving in depth, leading to slightly curved paths. While the model accurately predicts the initial lateral and depth landing positions, actors opt for a non-linear path towards this estimate. This decision might result from varying levels of uncertainty in lateral and depth estimates (compare panels B and C in Fig. 3) and likely differing associated displacement costs. Consequently, the actor independently utilizes running estimates in these dimensions at each time step to control velocity components. Due to lower variability in the lateral dimension, lateral movement progresses more rapidly, requiring fewer variable steps over time. This behavior aligns with the notion that interception errors are well explained by perceptual errors (de la Malla et al., 2018). Our explanation for the observed curved path differs from heuristics that assume outfielders consistently track the ball to generate curvatures (Fink et al., 2009; McBeath et al., 1995). In contrast, our model does not impose a specific gaze behavior, except for a minimal sampling rate required to update predictions. This allows a ball game player to divert her gaze to look somewhere else (e.g., acknowledging teammates’ positions) or use it in a flexible way (López-Moliner and Brenner, 2016). Fig. S1 in the SI shows individual paths followed by our participants. In this figure, we show that the spatial trajectories for the individual trials where the gaze is eventually diverted from the ball are not different from the others. However, our participants chose to track the ball almost continuously (Postma et al., 2014) (see Fig. S9 in SI) rather than predicting the interception location and moving in a straight line towards the interception location without looking at it. Continuous tracking of the ball allows the outfielder to cope with deviations from the expected trajectory (Brenner and Smeets, 2017; Fink et al., 2009) or minimize prediction uncertainty (Fooken et al., 2016; Mann et al., 2019).

Through initial estimates of landing positions and flight time, we offer a potential solution to the challenge of determining ball catchability (Postma et al., 2018). For instance, a rapidly increasing elevation angle combined with a very short time-to-contact *T*_*c*_ might indicate a ball that is coming fast and close, requiring swift action. In contrast, a slowly changing elevation angle with a large *T*_*c*_ might suggest that the ball is out of reach or not arriving for a significant duration. By bridging perceptual experience with objective motion dynamics, our model provides a concise yet robust framework to be tested in the future for understanding when a ball is deemed catchable.

The proposed model bridges the gap between the performance in ball game behavior and a more general perspective in visual perception which elucidates how 3D structure can be inferred from retinal motion given some rigidity assumptions (Longuet-Higgins and Prazdny, 1980; Ullman, 1979). In our case, participants can recover important 3D information (e.g. landing points) from the dynamics of optical variables under assumptions of size and gravity. Importantly, the model can cope with uncertainty of perceptual measurements in novel settings such as altered gravitational fields as evidenced by our study. This adaptability enables the model to make predictions under varying conditions, offering a nuanced solution to the long-standing outfielder problem.

## Methods

### Participants

We tested 12 participants (six self-identified women and six self-identified men). One participant had to be discarded due to a particularly noisy eye-tracker’s data (the filtering procedure removed more than 10% of the trials). Participants’ ages were between 22 and 33 years with normal or corrected-to-normal vision. All the participants were naïve to the experimental goals and volunteered to participate in the experiment. This study is part of an ongoing research program approved by the local ethics committee of the University of Barcelona in accordance with the Code of Ethics of the World Medical Association (Declaration of Helsinki).

### Apparatus

They performed the task by wearing a head-mounted display holding a controller with their dominant hand (all were right-handed). The experiment was performed on an Intel i7-based PC (Intel, Santa Clara, CA, USA)(i7-9700F). The stimuli were rendered using an NVIDIA GeForce (RTX 2060 SUPER) and sent to a wireless HTC Vive Pro head-mounted display (HMD) at 90 Hz per eye. Eye movements were recorded using a built-in eye tracker (Tobii Technology, 2011) sampled at 90 Hz.

### Stimuli

We used 10 different trajectory angles (Fig 1B). The ball’s initial position (x=0, z=40) was 40 m from the observer and laterally aligned with the observer’s initial position (lateral x=0, depth z=0). The interception location was always located 3 m away from the observer’s starting position describing horizontal trajectory angles of ±30, 60, 90, 120, 150 degrees (negative denotes left side and 90 deg corresponds to z=0) with respect to the observer’s initial location (see Fig. 1 B). We used both the standard gravitational acceleration (9.807 m/s2) at sea level and soccer ball size (0.22 m diameter) ± 10% of their respective standard values. In total, we had 90 (10 trajectories × 3 G × 3 size) conditions.

The ball’s initial height was vertically aligned to eye height on a trial-by-trial basis to account for any HMD slip and postural changes. Flying time was randomly selected from a uniform distribution ranging from 3.15 to 3.85 seconds (± 10% of 3.5 seconds). The initial and final positions, flight duration, and gravitational acceleration fully determine the initial vertical and horizontal velocities of the ball. Air resistance and other complex effects were neglected.

### Procedure

Prior to the experimental procedure, the participant and the experimenter tossed a standard-sized soccer ball (diameter 22 cm) back and forth to develop familiarity with the ball’s size. Each participant underwent a total of 10 blocks of 90 trials each. Each block was presented with one repetition of all combinations of gravity, size, and trajectory. The task was self-paced. Each block lasted for about 10-16 minutes. Participants completed 20 training trials before the main experimental procedure to familiarize themselves with the task and VR environment. The eye tracker was calibrated before each block. Calibration accuracy was tested with a custom program and always remained below 1.89 degrees error. Each trial was conducted as follows: (1) The participant was instructed to align both body and gaze while looking at the ball. Once aligned, the participant launched the ball by pressing a button. (2) Once the ball was in the air, the observers followed the ball visually while moving towards the interception point. (3) After completing 90% of the flight time, the ball was occluded and was no longer visible. The participants were instructed to head towards the position where they would have headed the ball and press a button to estimate the time-to-contact. Participants did not receive any feedback on their performance.

### Data analysis

The data obtained in this experiment were analyzed using R language (R Core Team, 2022). Gaze was categorized as being on the ball if the absolute vertical distance between the ball and gaze was lower than 6.5 degrees (Postma et al., 2014). The probability of gaze being on the ball was, on average, larger than 90% during the entire time the ball remained visible (see Fig. S1 in the SI). Since the direction of the ball (left or right) did not affect the temporal errors committed by our participants (t(10) = -0.05, p = .961) and our observers’ average heading angle (t(10) = -0.953, p = .363), we collapsed the results for the right and left trajectories for further analyses. For the final analysis, we removed those trials in which the frame rate was inconsistent, that is, the mean frame rate was lower than 81 fps. In addition, we removed trials in which the eye was detected by the eye tracker in less than 90% of the flight time and trials where the participant did not look at the ball at all (absolute average vertical distance between the ball and gaze was larger than 15 degrees). Finally, we excluded trials in which the response time was longer than 5 s. This procedure eliminated 265 trials (2.67% of the total).

### Locomotion

To analyze the paths traveled across trajectories, gravities, and sizes, we first normalized the paths based on the percentage of the distance covered. Each trial was then divided into a hundred steps, with the 100th step corresponding to the moment the participant pressed the trigger, indicating that the ball returned to eye height. We then computed the average heading angle for each participant, trajectory, gravitation, ball size, and step number. The heading is defined as the angular measure between the direction the participant is moving and ball’s initial position (0 degs indicates movement along the direct line between the starting position of the participant and that of the ball). For the ANOVA analysis, we aggregated the heading data by subject, trajectory, gravity, and ball size, focusing on the steps before occlusion and once the actor began moving. Prior to the analysis, we visually inspected the densities to check for a normal distribution of the angles. In this analysis, participants were treated as a random effect, while trajectory, gravity, and ball size were treated as fixed effects.

### The Model

#### Previous definitions

Fig. 1 A illustrates the general case of parabolic movement, with the observer’s initial position at the origin of the coordinate system, given by 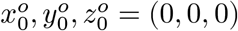. Assume that the observer’s eye-height is at plane *y* = 0. At time *t* = 0, a ball is launched from the point (*x*_0_, 0, *z*_0_) at eye height. The ball is seen at time *t* = *m* in the figure, and returns to eye height at time *t* = *T*, located at point (*x*_*T*_, 0, *z*_*T*_).

- *s* physical diameter of the target
- (*x*_*m*_, *y*_*m*_, *z*_*m*_) position of the ball at time *m*
- *υ*_0*y*_ Initial vertical component of the ball motion
- *υ*_*y*_, *υ*_*x*_, *υ*_*z*_ Vertical, lateral and depth components of parabolic motion (see Fig. 1 A)
- *υ*_*r*_ Radial component of the parabolic motion from the observer point of view (see Fig. 1 A)
- *υ*_*t*_ Tangential component of the parabolic motion from the perspective of the observer (normal to the radial component)
- *d* distance of the ball to the observer
- *T* is the total flight time (i.e. ball is above eye-height) and is given by: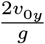
- *γ* elevation angle
- *θ* angle subtended by the ball
- *T*_*c*_ remaining time for the target to return at eye-height after movement onset at some specific position or moment of time.

Target position, angular variables (e.g. *θ, γ, β*), their temporal derivatives and remaining *T*_*c*_ are time dependent variables, but we will drop time indexes for simplicity. The height *y* of the ball at time *t* is given by (using *y*_0_ = 0):

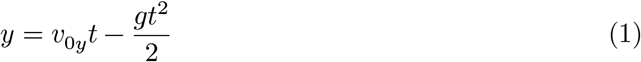

We will further assume that the depth position (*z*) of the ball is given by:

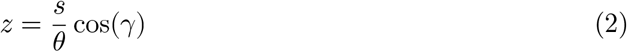

The tangent of the elevation angle *γ* at time *t* is:

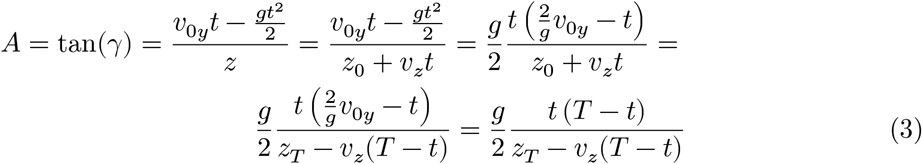

Where we have used: *z*_*T*_ = *z*_0_ + *υ* _*z*_*T* . From (3), we have:

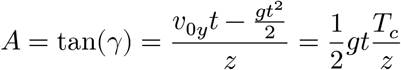

Therefore, when *t* ≠ 0,

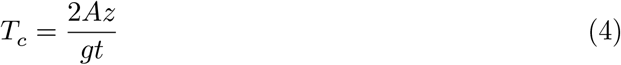

where *T*_*c*_ is the remaining time-to-contact (i.e. the ball returns at eye-height) at time *t*.

#### Estimation of *T*_*c*_, and *x*_*T*_ when *z*_*T*_ = 0

In this scenario, the ball final position is at the same depth as the initial position of the observer *z*_*T*_ = 0. Since *z* > 0, *υ*_*z*_ < 0 and *z*_*T*_ = 0, according to equation (3) we have that the tangent of *γ* is:

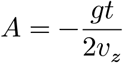

therefore,

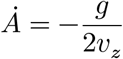

Furthermore, when the target is at position *z*, the remaining time is 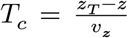 and since *z*_*T*_ = 0 we have:

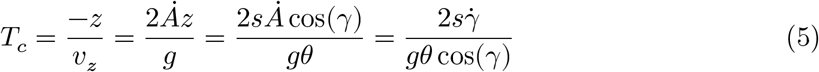

Equation 5 defines the remaining flight time (*T*_*c*_) as a function of optical variables, gravitational acceleration *g* and physical size *s*. Interestingly, this equation corresponds to the case when the ball falls on the initial location of the observer (Gómez and López-Moliner, 2013).

As for *x*_*T*_ (i.e., the final lateral position), we can calculate it using the formula:

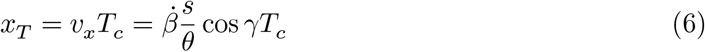

Substituting *T*_*c*_ in equation 6 with equation 5 we finally have:

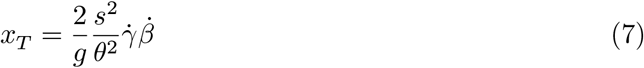

The final lateral position is fully specified by optical variables and *g* and *s*.

#### Estimation of *T*_*c*_, *x*_*T*_ and *z*_*T*_ when *t* = 0

In this case, when Δ*t* tends to 0, we can arrive at:

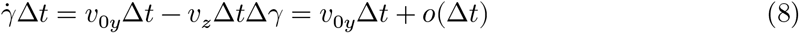

where *o* is the little *o* of Landau, so we can have:

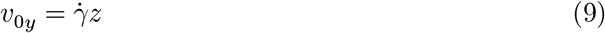

Therefore,

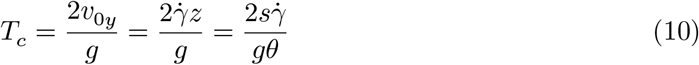

Since cos *γ* at *t* = 0 is 1, equations (10) and (5) are equivalent.

With respect to *z*_*T*_, we have that:

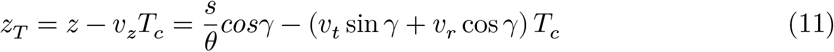

We have decomposed the velocity component in depth *υ* _*z*_ into the tangent *υ*_*t*_ and radial component *υ*_*r*_ (see Fig. 1 A). Since 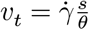 and 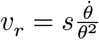 we can obtain *z*_*T*_ once we know *T*_*c*_:

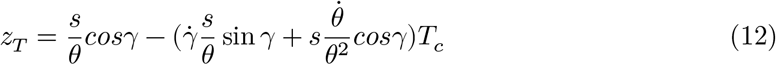

The final position in depth is specified at the initial moment by optical variables and *g* and *s* constants. The limiting factor in estimating *z*_*T*_ is the rate of expansion 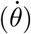. In our simulations, we introduced noise values for 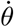 that exceeded known reported thresholds of 11% (Regan and Hamstra, 1993). Despite this, estimates of *z*_*T*_ remained robust. This robustness can be attributed not only to the dependence on *υ*_*r*_ (which is influenced by 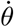) contribution from *υ*_*t*_.

In relation to *υ*_*x*_, like before, it is easy to solve it (see Fig. 1 A):

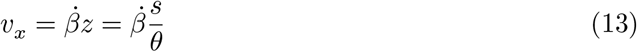

therefore,

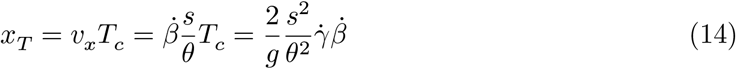

we obtain the same expression as in equation 7.

### Controller dynamics

The dynamics of the controller were inspired by the controller put forward in Fink et al. (2009). The radial and tangential accelerations are controlled respectively as follows:

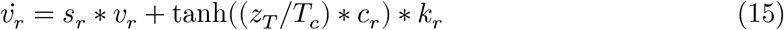

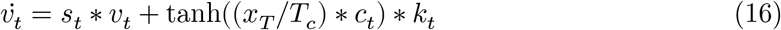

where *s*_*r*_ and *s*_*t*_ are damping terms and *K*_*r*_ and *K*_*t*_ are stiffness terms; *c*_*r*_ and *c*_*t*_ are thresholds terms that approximate a sigmoidal response to the control estimates. The values of *s, K* and *c* were obtained with the *optim* function implemented in the *stats* package of the R software. The optimization procedure minimized the negative log likelihood (nll) between the model predictions and the averaged (x, z) data points in our dataset. The optimization procedure was applied to trajectories in the 1G condition only. The values 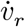 and 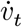 were integrated at each frame to update both velocity components of the actor (see R code in file “2_Controller.qmd”; Δ*t* = 0.05 s.). The resulting fitted parameters were: *s*_*r*_ = 0.735; *K*_*r*_ = 1.032; *c*_*r*_ = -0.565; *s*_*t*_ = 0.494; *K*_*t*_ = 1.545; *c*_*t*_ = -0.698.

After this optimization procedure, we run another optimization procedure minimizing the nll to obtain the weight of the Gravitation prior employing averaged data across the 3 gravitations and the 5 possible trajectory angles. The fitted value corresponded with a prior weight of 0.192 (*w* = 0.192).

Additionally, we implemented a second controller heuristic controller to fit our data. The implementation follows a similar approach, with some differences. In this case, the simulated outfielder’s movements were guided by linking them to the optical variables used in Fink’s (Fink et al., 2009) approach. Like their implementation, the jerk in the radial and tangential components of the movement relied on the first and second derivatives over time of tan(*ϕ*) and tan(*γ*), respectively:

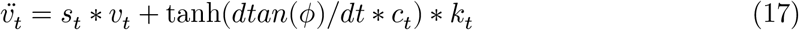

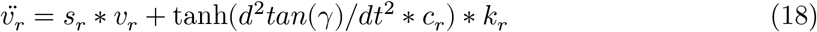

Similarly, *s*_*r*_ and *s*_*t*_ are damping terms and *K*_*r*_ and *K*_*t*_ are stiffness terms; *c*_*r*_ and *c*_*t*_ are thresholds terms that approximate a sigmoidal response to the control estimates. The values of *s, K* and *c* were obtained as before by minimizing the nll between simulated model trajectories and averaged (x, z) data points in our dataset. The values 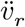 and 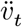 were integrated at each frame to update both actor velocity components (see R code in file “2_Controller.qmd”). We initially employed this procedure to estimate the values of *s*_*r*_, *s*_*t*_, *K*_*r*_, *K*_*t*_, *c*_*r*_, and *c*_*t*_ in the 1G condition with the resulting fitted parameters: *s*_*r*_ = 5.907; *K*_*r*_ = 2.797; *c*_*r*_ = 39.592.; *s*_*t*_ = 7.289; *K*_*t*_ = 13.998; *c*_*t*_ = 17.134. Subsequently, we applied this set of parameters to test the model’s performance in the gravity conditions other than 1G. However, since the predictive model incorporates an additional parameter (*w*) when tested in gravities different than 1G, we conducted further fitting procedures, as described in the Supplementary Information (SI), to enhance the flexibility of the heuristic model. These additional fitting procedures involved adjusting different controller terms in conditions other than 1G to achieve a better fit to the data.

## Acknowledgments

This work was funded by Grant Ref. PID2020-114713GB-I00 to JLM by MCIN/AEI/10.13039/501100011033. BA was supported by the fellowship FPU17/01248 from Ministerio de Educación y Formación Profesional of the Spanish government.

We would like to thank the late José Gómez for his considerable help that made this work possible. Also, we would like to thank Loes van Dam, Kai Streiling, Celine Honekamp and Cristina de la Malla for their constructive criticisms in previous versions of the manuscript.

