## Supplementary Information for "The predictive outfielder: a critical test across gravities"

Borja Aguado (1,2) and Joan López-Moliner (1)

1: Vision and Control of Action (VISCA) Group, Department of Cognition, Development and Psychology of Education, Institut de Neurociències, Universitat de Barcelona, Passeig de la Vall d'Hebron 171, 08035 Barcelona, Catalonia, Spain

2: Sensorimotor Control and Learning group, Centre for Cognitive Science, Department of Human Sciences, Institute for Psychology / Centre for Cognitive Science, Technische Universitat Darmstadt, Germany

#### SI 1: Individual trajectories per angle and participant

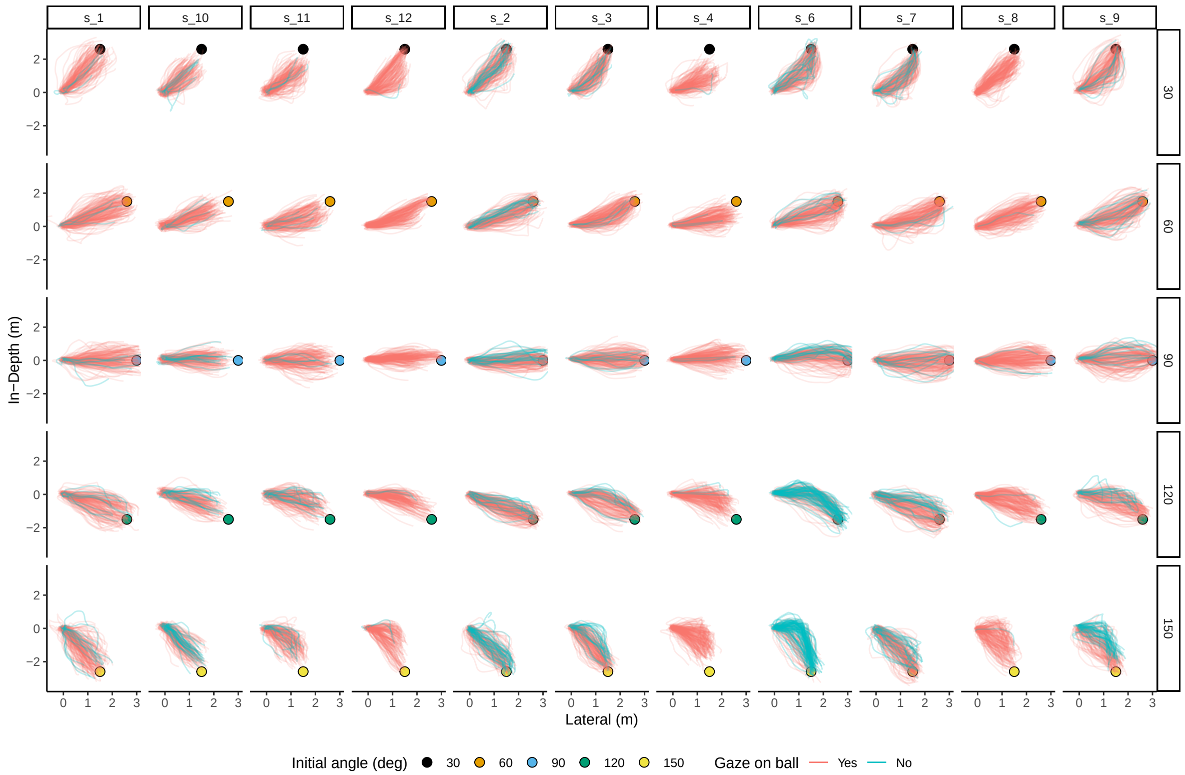


Fig. S 1: **Individual Trajectories by Trajectory Angle and Participant**. This figure is organized by trajectory angle (rows) and participant (columns). The color coding indicates whether the observer's gaze deviated more than 6.5 degrees on average from the ball during its trajectory, following the criteria of Postma et al. (2014). Colored dots mark the ball's landing location.

### SI 2: Testing for the use of initial vertical velocity and other variables in locomotion

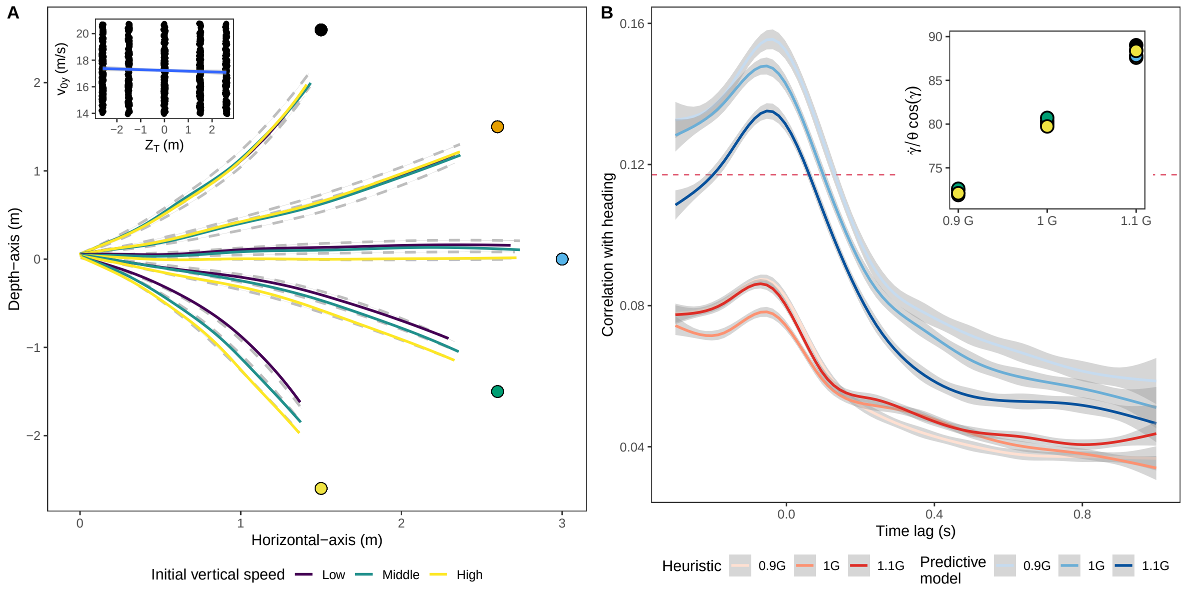


Fig. S 2: **Ruling Out Alternative Information Sources**. (A) Given the mathematical relationship among initial vertical velocity $v_{0y}$, flight time $T_c$, and gravity (G), expressed as $T_c = 2v_{0y}/G$, it is crucial to discount the possibility that differences in trajectories across gravities are due to $v_{0y}$. We categorize participant trajectories by $v_{0y}$ (3 levels) with the average difference between the highest and lowest levels exceeding 10% for all trajectories. This suggests that the lack of trajectory differences for balls landing in front of the observer is not due to discrimination difficulty. Dashed grey lines represent paths under different gravities (as shown in Fig. 2 of the article), with colored dots marking the ball's landing points. The inset confirms the absence of a correlation between final in-depth position and $v_{0y}$, indicating that the latter does not predict in-depth location. (B) The cross-correlation function between heading and predictive control variables ($Z_T/T_C$ and $X_T/T_C$), calculated as $\sqrt{(Z_T/T_C)^2 + (X_T/T_C)^2}$, is shown in varying shades of blue for different gravity values. In red, we depict the correlation between heading and heuristic optical variables ($d^2 \tan\gamma$ and $d \tan\phi$) controlling in-depth and lateral movement, respectively, calculated as $\sqrt{(d^2 \tan\gamma)^2 + (d \tan\phi)^2}$. Correlations above the red dotted line are significantly larger than zero.

#### SI 3: Effect of different Earth gravity weights on the path followed towards the ending point

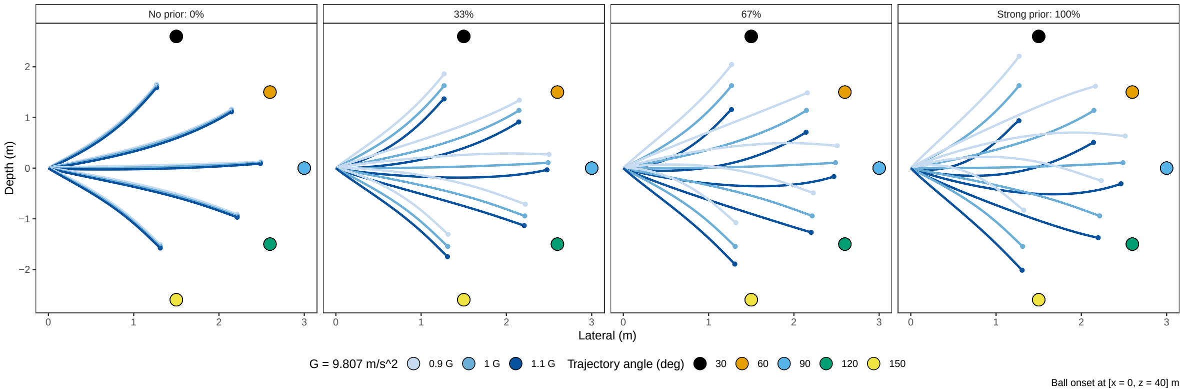


Fig. S 3: **Paths of a Simulated Predictive Agent Across Various Earth Gravity Weights**. Consistent with our model and the agent's controller implementation, an increase in the weight (w) given to Earth's gravity (1G), relative to the weight (1-w) for the simulated gravity affecting the ball's motion, results in a more distinct separation of trajectories across different gravities (different panels). The optimal Earth gravity weight for the best fit is found to be 19.2%.

### SI 4: Additional flexibility to heuristic models (i)

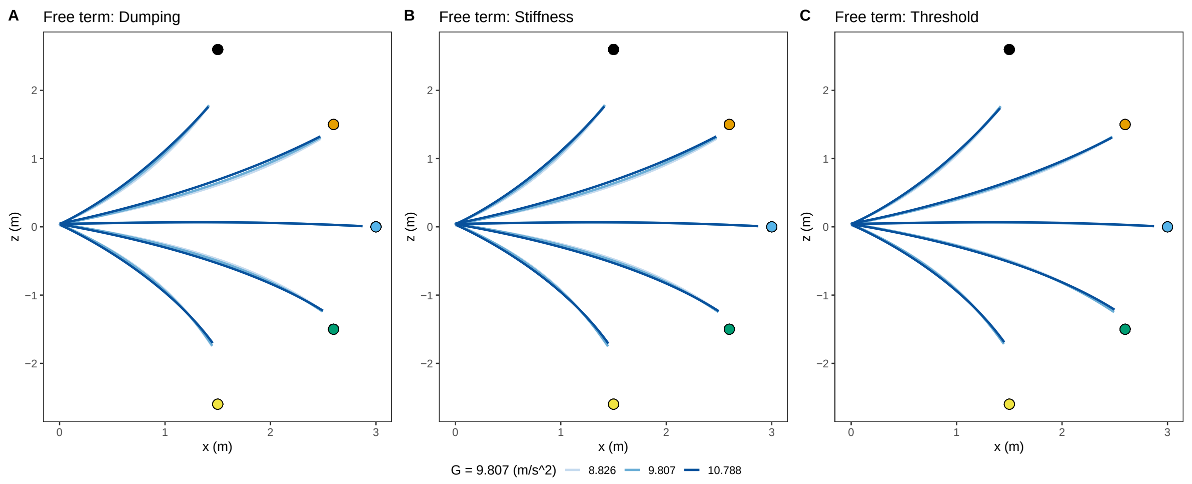


Fig. S 4: **Trajectories predicted by the heuristic strategy freeing different parameters**. Fig. 2B in the article presents the GOAC heuristic model's predictions, showing a symmetrical pattern in trajectories landing in front of and behind the observer for the same gravity values. This symmetry, stemming from the GOAC model's aim to cancel perceived optical acceleration, contrasts with empirical paths that do not follow this symmetry, highlighting the model's limitations. Fig. 2C, in contrast, shows how the predictive controller, incorporating gravity knowledge, successfully replicates empirical paths. Using six parameters fitted in the 1G condition and an additional weight parameter (w=0.19) for other gravity conditions, the predictive model accounts for trajectory divergence across different gravities, totaling seven parameters. We further explored the heuristic model's flexibility by allowing additional parameters to vary independently (different panels) in the 0.9G and 1.1G conditions, leading to a total of 10 parameters (6 for G=1, 2 each for 0.9G and 1.1G). The fitting code is available in '2_Controller.qmd' at https://osf.io/bcp28/?view_only=e8cfd4e604304cbab630903eead874d6. Despite these modifications, the heuristic strategy fails to predict empirical data patterns. The figure displays various parameter adjustments but does not align with the observed empirical deviations.

#### SI 5: Additional flexibility to heuristic models (ii)

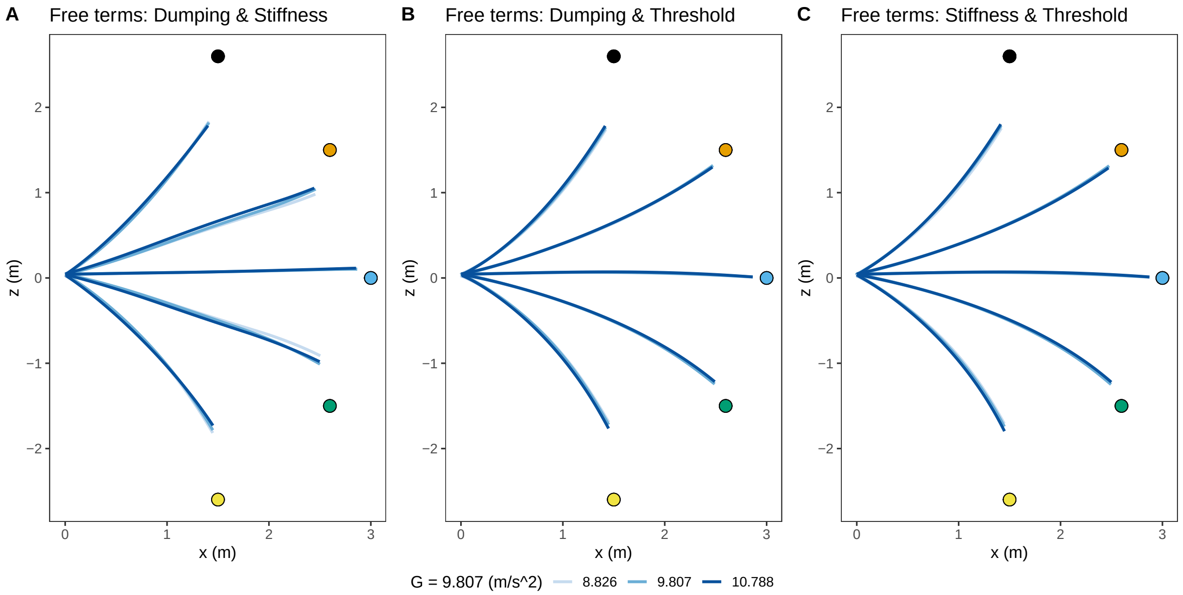


Fig. S 5: Trajectories predicted by the heuristic strategy freeing four parameters (two per each term -e.g. dumping- of the controller) in the conditions different than 1G. To introduce additional flexibility, albeit at the expense of increasing the number of free parameters, we performed the same procedure again. This time, we allowed two terms, such as damping and stiffness (a total of four parameters), to vary independently in the 0.9G and 1.1G conditions: 6 parameters (G=1) + 4 parameter (0.9G) + 4 parameter (1.1G) = 14 parameters in total.

#### Model comparisons

Table 1: Summary of model parameters, log likelihood, AIC and probability of minimizing information loss relative to the predictive model.

| Model | Num pars | Log Likelihood | AIC | Prob[1] |
|---|---|---|---|---|
| Predictive | 7 | -9411.889 | 18837.78 | ----------- |
| GOAC | 6 | -9452.342 | 18916.68 | 7.356003e-18 |
| Dumping free | 10 | -9449.046 | 18918.09 | 3.631951e-18 |
| Stiffness free | 10 | -9448.005 | 18916.01 | 1.028402e-17 |
| Threshold free | 10 | -9447.784 | 18915.57 | 1.283099e-17 |
| Dumping+Stiffness | 14 | -9430.621 | 18881.24 | 3.645979e-10 |
| Dumping+Threshold | 14 | -9444.685 | 18917.37 | 5.207862e-18 |
| Stiffness+Threshold | 14 | -9447.221 | 18922.44 | 4.123366e-19 |

[1] exp((aic_pred-aic_model)/2) times as probable as the predictive model to minimize the information loss.

While increasing flexibility enhances the fit of the heuristic model, the best-performing heuristic model, obtained by allowing both the damping term and stiffness to vary, still falls short of matching the predictive model. There is no heuristic model that demonstrates an equivalent likelihood to the predictive model in terms of minimizing information loss as shown in the last column (prob) of the table.

Despite introducing added flexibility by allowing different parameters for each gravity condition, the heuristic model still failed to create distinctly separated paths in the correct direction within trajectory angles. This outcome suggests that the heuristic model strives to find a compromise or a set of parameters that can provide a reasonable fit for both types of trajectories within a single gravity condition. However, this approach seems to generate a mean path that lacks the fidelity to capture the variations present in the experimental data. It's worth noting that the predictive model effortlessly reproduces the desired separation in the correct direction using the same parameter (w) across all gravities and trajectory angles.

#### SI 6: Error map temporal predictions

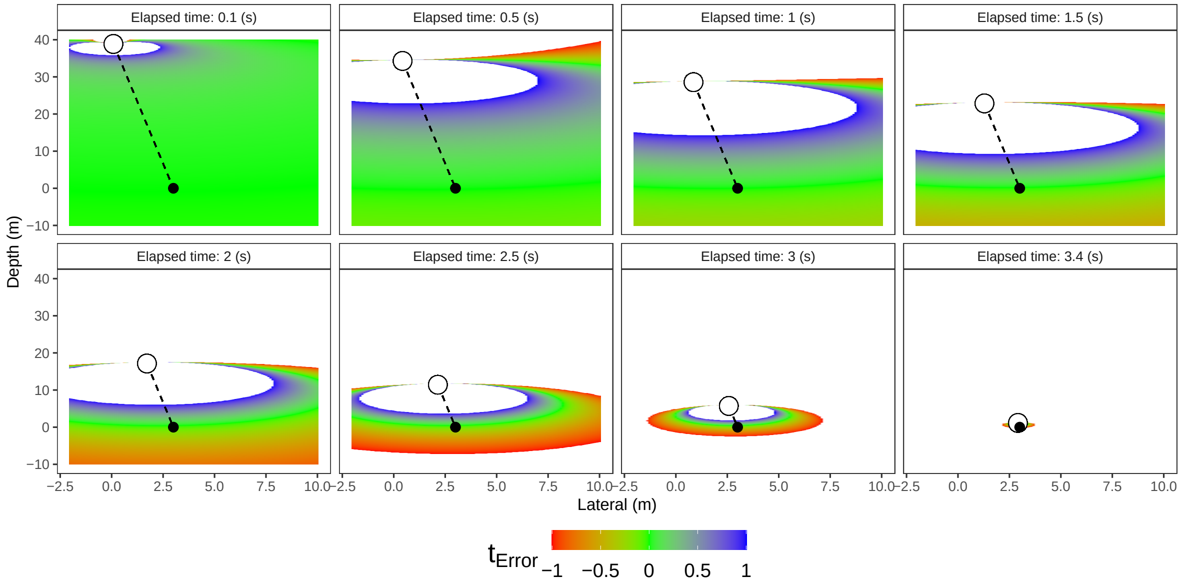


Fig. S 6: Temporal error map based on the model estimate of remaining time-to-contact ($T_c$) at different moments (panels). We plot temporal errors at various time intervals following the initiation of the ball's motion (with white and black dots marking the ball's initial and landing positions, respectively). The color-coded representation in the (x, z) plane illustrates the temporal error linked to the model's estimate when the observer is positioned at that particular (x, z) coordinate. As can be seen, our predictive model accurately estimates the remaining time-to-contact at the onset of motion irrespective of the observer's viewing point (first panel). This initial accuracy enable the observer to gauge the necessary speed to reach the interception point. An animated version (GIF) of the figure can be obtained in the 3_Temporal_Predictions_GIF.R file in the OSF. The white and the black points represent the ball and the ending point respectively. White areas correspond with temporal errors larger than 1 second. Those errors are not represented to increase the resolution of of the color scale.

#### SI 7: Relevant optic variables used by heuristic strategies

Previous heuristic strategies assume that the outfielder would control the navigation by using error-nulling tactics based on different kinds of optic information (1–3)

In the figure below, the reader can see how three strategies: OAC (Fig S7 A), GOAC (Fig S7 A and B) and LOT (Fig S7 C) would be congruent with the trajectory followed by our predictive agent when catching a ball in the 1G condition.

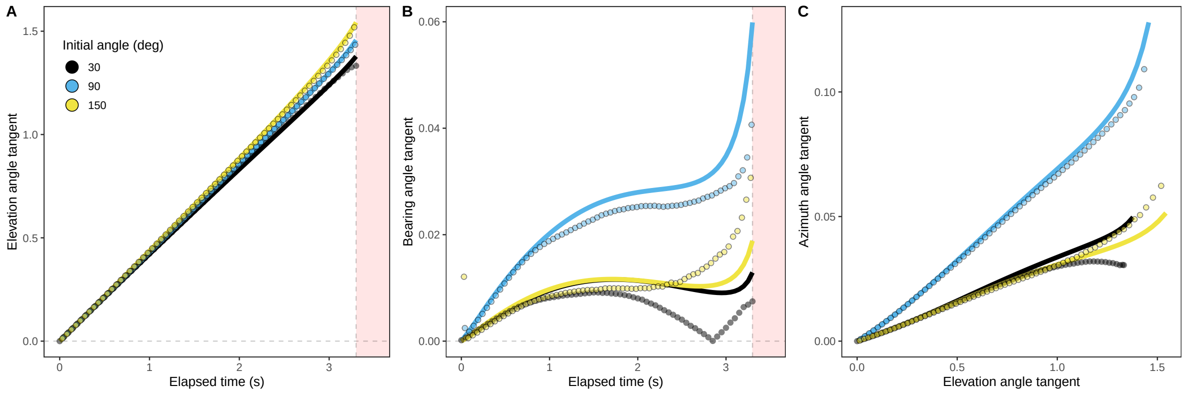


Fig. S 7: In this figure, lines indicate the predictions of the simulated predictive agent and dots indicate the corresponding optical information based on the empirical trajectories. (A) Tangent of the elevation angle ($\gamma$) versus the elapsed time. In this case, the elevation angle increases linearly over the course of the trajectory. (B) Tangent of the bearing angle ($\phi$) versus the elapsed time. In this case, the bearing angle remains relatively constant after 1 sec and during a large part of the trajectory. (C) Tangent of the bearing angle versus the tangent of the elevation angle. As predicted by the LOT, this ratio increases approximately linearly during the course of the trajectory.

#### SI 8: Predictive control variable

Analysis of our data reveals that participants often initiate movement quite early in the trajectory of the ball. While this does not conclusively confirm a predictive approach, it suggests that participants aren't merely reacting to the ball's current state but may be estimating its future path based on early cues. This early initiation of movement would align more with a predictive framework.

In Fig S8 A, we observe that the depth velocity of our participants begins to diverge before the 0.5-second mark (shaded areas represent 95% confidence intervals). Statistically significant separations (non-overlapping 95%-CI) of these velocities occurs notably earlier than a 20% threshold difference of the rate of change of the elevation angle (maximum difference at 0.5 is 4.35%), as illustrated in Fig S8 B, which is consistent with previous findings (4). A difference of 20% is the minimum change of speed reported (5) to detect acceleration.

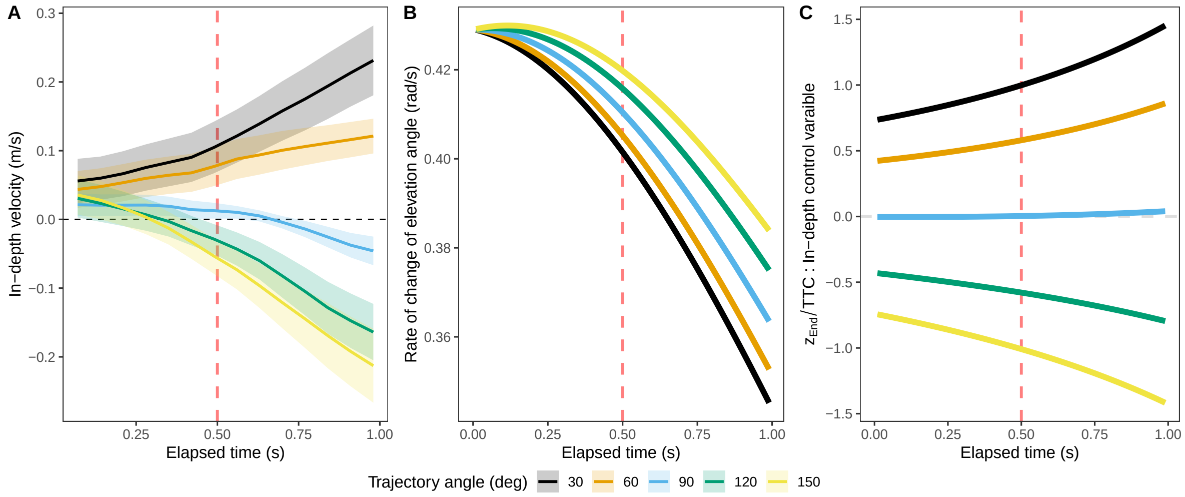


Fig. S 8: A) Average in-depth velocity for different trajectory angles (color code). Shaded area indicates a 95% confidence interval. B) Rate of change of the elevation angle ($\dot{\gamma}$). C) Variables estimated by the predictive actor to control the in-depth velocity: ratio of the in depth landing point to the remaining flight time: $Z_t/T_c$. This would be a cue for the necessary speed to start running.

#### SI 9: Gaze in ball

Our participants tracked the ball consistently across trials during the period of the trajectory where the ball was visible consistently with others findings (6).

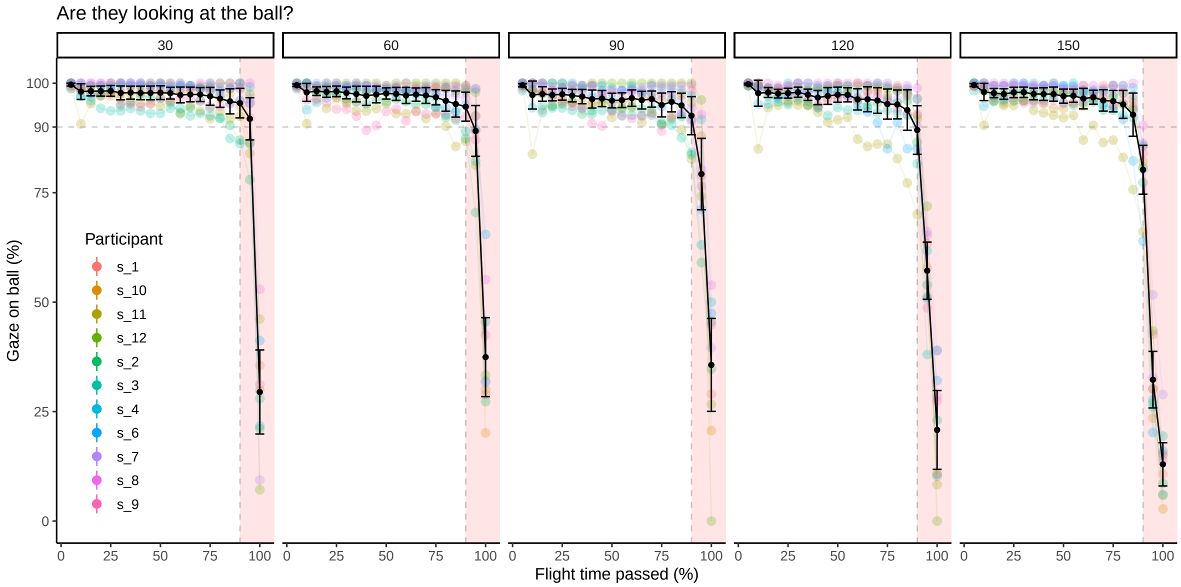


Fig. S 9: Percentage of gaze on ball (av. error gaze/ball < 6.5 deg) againts 20th of flight time passed (x-axis), per participant (color code) and trajectory (panels). The red shaded area indicates the portion of the trajectory when the ball was occluded from view.
